# Gene-Based Analysis in HRC Imputed Genome Wide Association Data Identifies Three Novel Genes For Alzheimer’s Disease

**DOI:** 10.1101/374876

**Authors:** Emily Baker, Rebecca Sims, Ganna Leonenko, Aura Frizzati, Janet Harwood, Detelina Grozeva, GERAD/PERADES, IGAP consortia, Kevin Morgan, Peter Passmore, Clive Holmes, John Powell, Carol Brayne, Michael Gill, Simon Mead, Reiner Heun, Paola Bossù, Gianfranco Spalletta, Alison Goate, Carlos Cruchaga, Cornelia van Duijn, Wolfgang Maier, Alfredo Ramirez, Lesley Jones, John Hardy, Dobril Ivanov, Matthew Hill, Peter Holmans, Nicholas Allen, Paul Morgan, Julie Williams, Valentina Escott-Price, GERAD Consortium, ADGC Consortium, CHARGE Consortium, EADI Consortium

## Abstract

A novel POLARIS gene-based analysis approach was employed to compute gene-based polygenic risk score (PRS) for all individuals in the latest HRC imputed GERAD (N cases=3,332 and N controls=9,832) data using the International Genomics of Alzheimer’s Project summary statistics (N cases=13,676 and N controls=27,322, excluding GERAD subjects) to identify the SNPs and weight their risk alleles for the PRS score. SNPs were assigned to known, protein coding genes using GENCODE (v19). SNPs are assigned using both 1) no window around the gene and 2) a window of 35kb upstream and 10kb downstream to include transcriptional regulatory elements. The overall association of a gene is determined using a logistic regression model, adjusting for population covariates.

Three novel gene-wide significant genes were determined from the POLARIS gene-based analysis using a gene window; *PPARGC1A, RORA* and *ZNF423*. The *ZNF423* gene resides in an Alzheimer’s disease (AD)-specific protein network which also includes other AD-related genes. The *PPARGC1A* gene has been linked to energy metabolism and the generation of amyloid beta plaques and the *RORA* has strong links with genes which are differentially expressed in the hippocampus. We also demonstrate no enrichment for genes in either loss of function intolerant or conserved noncoding sequence regions.

## Introduction

Late Onset Alzheimer’s disease (LOAD) is a devastating neurodegenerative condition with a significant genetic heritability (Gatz et al. [2006]). The *apolipoprotein E* (*APOE*) gene is the strongest genetic risk factor for LOAD (Strittmatter et al. [1993]). Subsequently, more genes were found to be associated with AD development. The Genetic and Environmental Risk in Alzheimer’s Disease Consortium (GERAD) published a Genome-Wide Association Study (GWAS) that identified novel variants in *CLU* and *PICALM* which were associated with AD (Harold et al. [2009]). Concurrently, the European Alzheimer’s Disease Initiative (EADI) identified the *CR1* and *CLU* loci to associate with AD (Lambert et al. [2013]). Subsequent publications by GERAD, the Alzheimer’s Disease Genetic Consortium (ADGC) and Cohorts for Heart and Aging Research in Genomic Epidemiology (CHARGE) Consortium identified a further 5 novel loci (Hollingworth et al. [2011], Naj et al. [2011], Seshadri et al. [2010]). The International Genomics of Alzheimer’s Project (IGAP) (Lambert et al. [2013]) consortium is an amalgamation of these four different genetic groups (GERAD, EADI, ADGC and CHARGE). Meta-analysis of the 4 GWAS datasets determined 11 novel variants associated with AD were determined. A gene-based analysis has been undertaken in the IGAP AD data using Brown’s method (Brown [1975]). This approach determined two additional novel genes; *TP53INP1* and *IGHV1-67* (Escott-Price et al. [2014]). Additionally, low frequency risk variants have been identified through next generation sequencing (*TREM2*) (Guerreiro et al. [2013]) and an whole-exome association study (*PLCG2, TREM2* and *ABI3* (Sims et al. [2017])).

Set-based analysis is an alternative to Genome-Wide Association Study (GWAS) analyses, which considers the association of an individual Single Nucleotide Polymorphism (SNP) with disease. Set-based analyses provide more power due to the aggregate effect of multiple SNPs being larger than that of individual SNPs. For example, determining the association of genes rather than SNPs, is beneficial since genes are more robust across different populations (Li et al. [2011]). Set-based analyses are being widely used in the field and as expected, are able to identify novel genes or pathways associated with disease. Pathways clustering in eight areas of biology have been found to be associated with AD using the ALIGATOR (Holmans et al. [2009]) algorithm (Jones et al. [2010]) (Jones et al. [2015]). Additionally, 134 gene-sets have been identified as being associated with Schizophrenia (SZ) that are related to nervous system function and development, where gene-sets are defined from single gene functional studies (Pocklington et al. [2015]). Other gene and gene-set analyses were considered in a SZ study investigating exomic variation which determined an enrichment in genes whose messenger ribonucleic acid (mRNA) binds to fragile X mental retardation protein (FMRP) and Loss of Function (LoF) intolerant genes (Leonenko et al. [2017]).

The aim of the current study is to identify novel genes associated with AD using the largest up-to-date reference SNP panel, the most accurate imputation software and the latest published gene-based analysis approach. In this study, we used the GERAD data (Harold et al. [2009]) which have recently been imputed using the latest Haplotype Reference Consortium data (HRC). Polygenic Linkage disequilibrium-Adjusted Risk Score (POLARIS) (Baker et al. [2018]) is a powerful gene-based method which produces a risk score per person per gene, adjusts for Linkage Disequilibrium (LD) between SNPs and informs the analysis with summary statistics from an external data set. POLARIS, unlike standard Polygenic Risk Score (PRS) does not require data to be pruned for LD prior to analysis, so is able to incorporate information from a larger number of SNPs. We employed the POLARIS approach (Baker et al. [2018]) and using the individual genotypes in the GERAD imputed data, produced the risk score for each individual for every gene considered. The IGAP (Lambert et al. [2013]) SNP summary statistic data, where the individuals from GERAD study were excluded, were used to generate the gene-based PRS. Furthermore, previous studies (e.g. Pardinas et al. [2018]) have suggested that genes associated with SZ are evolutionary constrained, likely due to functional importance during the neurodevelopmental stages. Since AD is a post-reproductive disorder, we hypothesised and tested whether genes associated with AD reside in conserved noncoding sequences or whether there is an enrichment of genes which are LoF intolerant.

## Methods

The Haplotype Reference Consortium (HRC), version r1.1 2016, was used to impute GERAD genotype data on the Michigan Imputation Server (Das et al. [2016]), which to date, allows most accurate imputation of genetic variants. Imputed genotype probabilities (also known as dosages) were converted to the most probable genotype with a probability threshold of 0.9 or greater. SNPs were removed if: their imputation INFO-score<0.4, minor allele frequency (MAF)<0.01, missingness of genotypes≥0.05 or HWE<10^-6^. A total of 6,119,694 variants were retained. To correct for population structure and genotyping differences, all analyses were adjusted for gender and the top 15 principal components.

POLARIS was applied to this GERAD (3,332 cases, 9,832 controls) imputed data, using the IGAP (Lambert et al. [2013]) data (17,008 cases, 37,154 controls) excluding GERAD subjects (IGAPnoGERAD) as an external dataset to improve power. There were 3,169,839 SNPs in common between imputed GERAD and IGAP summary statistics data. The GERAD imputed data contain individual genotypes for every SNP, enabling the production of a risk score per person, and the IGAPnoGERAD data contains effect sizes for every SNP, which are used to weight the risk score. A gene-based risk score was produced for every individual in the GERAD data.

POLARIS adjusts for LD between SNPs and therefore, the SNPs were not pruned for LD and the entire data were used in this analysis. POLARIS adjusts for LD by using spectral decomposition of the correlation matrix between SNPs. Such a matrix was derived for each gene using the individual genotypes from the GERAD imputed data. To ensure that SNP alleles were coded in the same direction across both independent datasets; IGAPnoGERAD and imputed GERAD if alleles in IGAPnoGERAD were coded in the opposite direction to those in GERAD, the summary effect size for the SNP was inverted. SNPs with alleles AT, TA, CG or GC were excluded.

MAGMA (de Leeuw et al. [2015]) software offers an alternative approach to gene-based analyses. It can either be applied to summary statistics, or to the individual genotypes. We run both options of MAGMA software in our GERAD HRC imputed data. These analyses however cannot use additional (IGAPnoGERAD) information, but include all SNPs (N=6,119,694), as merging with the IGAPnoGERAD summary statistic data is not required.

SNPs were assigned to genes using GENCODE (v19) gene models (Harrow et al. [2012]). Only genes with known gene status and those marked as protein coding were used. Two different gene windows were considered, the first used no window around the gene, only SNPs within the start and end position of the gene were included, and the second considered SNPs which were within 35kb upstream and 10kb downstream of the gene. These windows were used since it is likely to contain transcriptional regulatory elements (Network and Consortium Pathway Analysis Subgroup of Psychiatric Genomics [2015]). SNPs which belong to multiple genes were assigned to all those genes. In the HRC imputed GERAD data, 1,122,570 SNPs were assigned to 17,072 distinct genes when no gene window was used and 2,296,690 SNPs were assigned to 18,087 genes when the gene window was used.

A POLARIS score was produced for each of these genes, and the overall association of the gene with AD is determined using a logistic regression model, adjusting for population covariates.

It was then assessed whether genes determined from the gene-based analysis were enriched in conserved regions; both for genes which are evolutionary constrained LoF intolerant and those which reside in conserved noncoding sequences. Genes which are evolutionary constrained were determined using the Exome Aggregation Consortium (ExAC) which contains high quality exome sequence data for 60,706 subjects (Lek et al. [2016]). This database contains a list of genes, so the number of genes from our analysis which reside in this list were determined. The conserved noncoding regions were defined from Babarinde et al. [2016], this data contains genomic locations for the conserved noncoding regions. Genes from our gene-based analysis which overlap these regions were determined. These regions are less likely to harbour variants of a strong effect or are less prone to variation (Babarinde et al. [2016]). The number of genes from the gene-based analysis with a p-value below either a nominal (0.05) or gene-wide threshold (2.5 × 10^-6^) and were in conserved regions were determined. The 2×2 contingency tables (not presented) have columns containing the number of genes with p-values above and below the p-value threshold and the rows show the number of genes in/out of conserved regions. The strength of association between genes residing in conserved regions and gene p-value was assessed using a chi-squared test (or Fisher’s exact, when the cell counts in the 2×2 table were small). Since the same SNPs may be assigned to multiple genes, it is not possible to assume independence between genes. Therefore, prior to the analysis, genes within 250kb of one another were removed, retaining the most significantly associated gene. However, including all genes regardless of LD (not presented) does not change the result.

## Results

For the imputed GERAD data, the SNPs are assigned to 17,072 genes. Of these genes, six reach gene-wide significance when no gene window is used. These genes are *CLU, BCL3, PVRL2, TOMM40, APOE* and *CLPTM1* (see details in Table I); these have all been previously identified as being associated with AD (Harold et al. [2009], Lambert et al. [2013]). The majority of these associations are influenced by *APOE* since these genes are close in location to *APOE*. These results are shown on the Manhattan plot in Figure 1, the gene-wide significant genes are shown in green, and those with a suggestive p-value (2.5 × 10^-6^ < p < 0.00001) are shown in blue; these are *DAB1, ZNF35* and *RORA*.

**Table I:** Gene-Wide Significant Genes from POLARIS Gene-based Analysis in AD Imputed Data

|  |  |  | POLARIS |  |  |
| --- | --- | --- | --- | --- | --- |
| Chr | Gene | No. of SNPs | Beta | SE | P-value |
| 8 | *CLU* | 25 | 0.521 | 0.1048 | $6.8 \times 10^{-7}$ |
| 19 | *BCL3* | 5 | 0.927 | 0.1499 | $6.1 \times 10^{-10}$ |
| 19 | *PVRL2* | 95 | 0.637 | 0.0532 | $5.7 \times 10^{-33}$ |
| 19 | *TOMM40* | 20 | 0.454 | 0.0351 | $2.9 \times 10^{-38}$ |
| 19 | *APOE* | 1 | 2.247 | 0.2679 | $4.9 \times 10^{-17}$ |
| 19 | *CLPTM1* | 93 | 0.461 | 0.0965 | $1.8 \times 10^{-6}$ |

**Figure 1:**
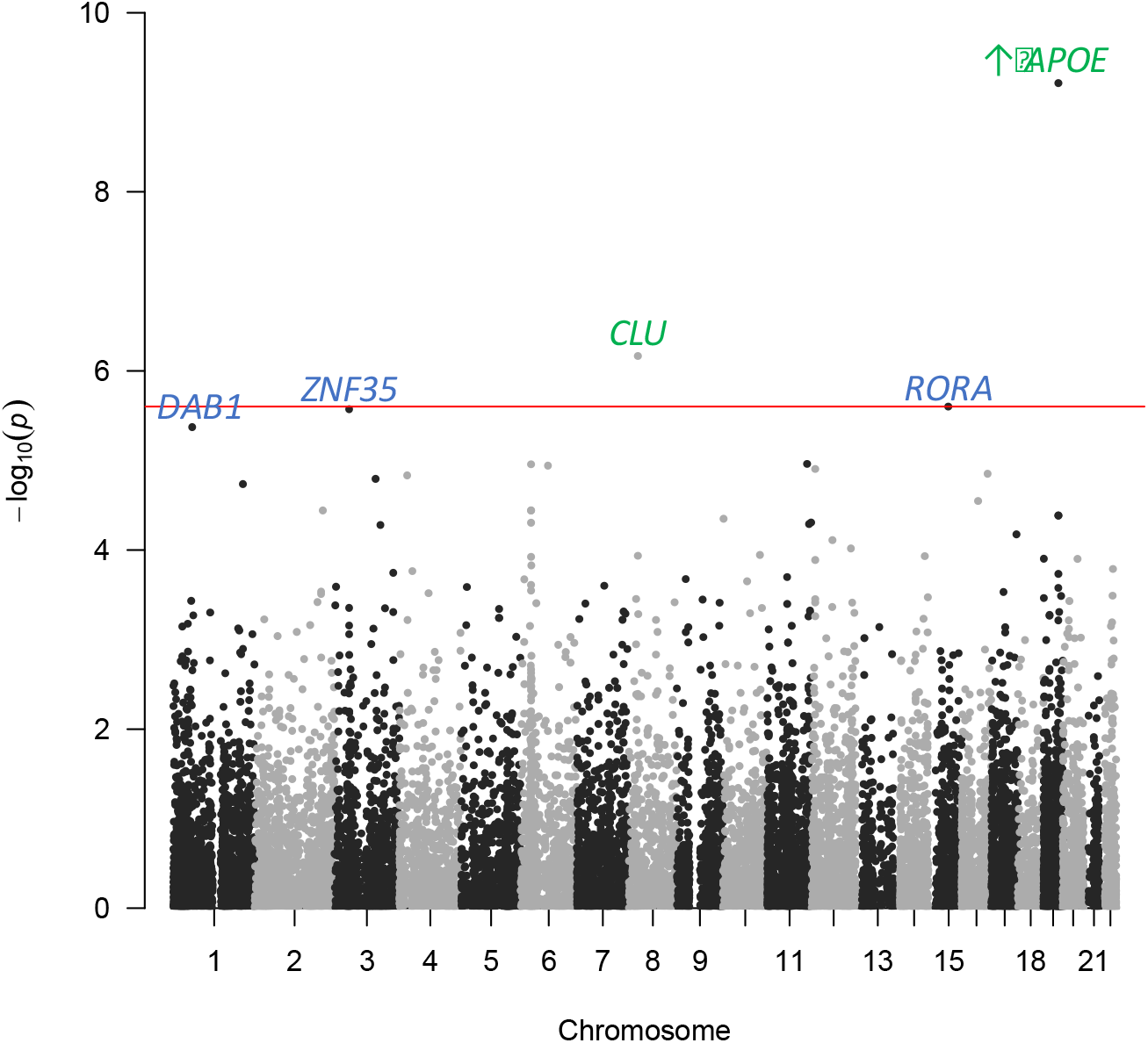
Manhattan Plot for the POLARIS Gene-Based Analysis in Imputed GERAD data

MAGMA can be applied to both individual genotype data (termed MAGMA-PCA) or to summary statistic data (termed MAGMA-SUMMARY). The gene-wide significant genes from the imputed GERAD data using MAGMA-PCA are in Table II. This method finds three genes below the gene-wide threshold, all of which are in the *APOE* region; *BCL3, PVRL2* and *TOMM40* (Harold et al. [2009], Lambert et al. [2013]).

**Table II:** Gene-Wide Significant Genes from MAGMA-PCA Gene-based Analysis in GERAD Imputed Data

| Chr | Gene | No. of SNPs | MAGMA-PCA P-value |
| --- | --- | --- | --- |
| 19 | *BCL3* | 5 | $2.7 \times 10^{-8}$ |
| 19 | *PVRL2* | 92 | $4.9 \times 10^{-35}$ |
| 19 | *TOMM40* | 20 | $1.6 \times 10^{-48}$ |

Table III shows the gene-wide significant results for the gene-based analysis in the imputed GERAD using MAGMA-SUMMARY. This method uses the summary statistics of the imputed GERAD data rather than the individual genotypes. This analysis determines four gene-wide significant genes, all of which have been previously identified or are part of the *APOE* region (Harold et al. [2009], Lambert et al. [2013]).

**Table III:** Gene-Wide Significant Genes from MAGMA-SUMM Gene-based Analysis in GERAD Imputed Data

| Chr | Gene | No. of SNPs | MAGMA-SUMM P-value |
| --- | --- | --- | --- |
| 8 | *CLU* | 25 | $1.5 \times 10^{-6}$ |
| 19 | *BCL3* | 5 | $9.5 \times 10^{-7}$ |
| 19 | *PVRL2* | 95 | $1.6 \times 10^{-13}$ |
| 19 | *TOMM40* | 20 | $4.0 \times 10^{-26}$ |

This analysis was repeated using a window around the gene of 35kb upstream and 10kb downstream since transcriptional regulatory elements are likely to be contained within this window (Network and Consortium Pathway Analysis Subgroup of Psychiatric Genomics [2015]). The gene-based results for the 18,087 genes are plotted on a Manhattan plot in Figure 2. The 12 gene-wide significant genes from this analysis are shown in Table IV. Again, a large number of genes reside on chromosome 19 and these are likely influenced by the large effect of *APOE*. Three novel genes have been identified from this analysis: *PPARGC1A, RORA* and *ZNF423*. The *PPARGC1A* gene has been linked to energy metabolism and the generation of amyloid beta plaques (Katsouri et al. [2016]) and has potential relevance to the human aging process. The *RORA* gene has strong links with genes which are differentially expressed in the hippocampus (Acquaah-Mensah et al. [2015]). The *ZNF423* gene resides in an AD-specific protein network which also includes other AD-related genes such as *APOE, CLU, ABCA7, TREM2* etc. (Hu et al. [2017]). In addition, the *SCARA3* gene overlaps *CLU* which has previously been identified as being associated with AD (Harold et al. [2009], Lambert et al. [2013]), and has additionally been found to be associated with total brain volume (Chauhan et al. [2015]). Note that the gene based analysis on IGAP data (Escott-Price et al. [2014]) identified some genes not found using this POLARIS approach. This could be because the POLARIS analysis focused on GERAD data rather than the IGAP data, and therefore does not have the same SNP coverage, or IGAP results could be influenced by other consortia.

**Table IV:** Gene-Wide Significant Genes from POLARIS Gene-based Analysis in GERAD Imputed Data Using a Gene Window (35kb Upstream and 10kb Downstream)

|  |  |  | POLARIS |  |  |
| --- | --- | --- | --- | --- | --- |
| Chr | Gene | No. of SNPs | Beta | SE | P-value |
| 4 | *PPARGC1A* | 480 | 0.877 | 0.1851 | $2.2 \times 10^{-6}$ |
| 8 | *SCARA3 (CLU)* | 240 | 0.526 | 0.1064 | $7.8 \times 10^{-7}$ |
| 15 | *RORA* | 1813 | 0.334 | 0.0674 | $7.4 \times 10^{-7}$ |
| 16 | *ZNF423* | 1056 | 0.551 | 0.1163 | $2.1 \times 10^{-6}$ |
| 19 | *BCL3* | 88 | 0.377 | 0.0674 | $4.2 \times 10^{-9}$ |
| 19 | *CBLC* | 50 | 0.605 | 0.1161 | $1.8 \times 10^{-7}$ |
| 19 | *BCAM* | 71 | 0.556 | 0.0543 | $1.4 \times 10^{-24}$ |
| 19 | *PVRL2* | 160 | 0.546 | 0.0299 | $9.4 \times 10^{-75}$ |
| 19 | *TOMM40* | 108 | 0.500 | 0.0298 | $3.4 \times 10^{-63}$ |
| 19 | *APOE* | 55 | 0.520 | 0.0315 | $4.4 \times 10^{-61}$ |
| 19 | *APOC1* | 34 | 0.475 | 0.0315 | $1.5 \times 10^{-51}$ |
| 19 | *APOC4-APOC2* | 62 | 0.615 | 0.0871 | $1.6 \times 10^{-12}$ |

**Figure 2:**
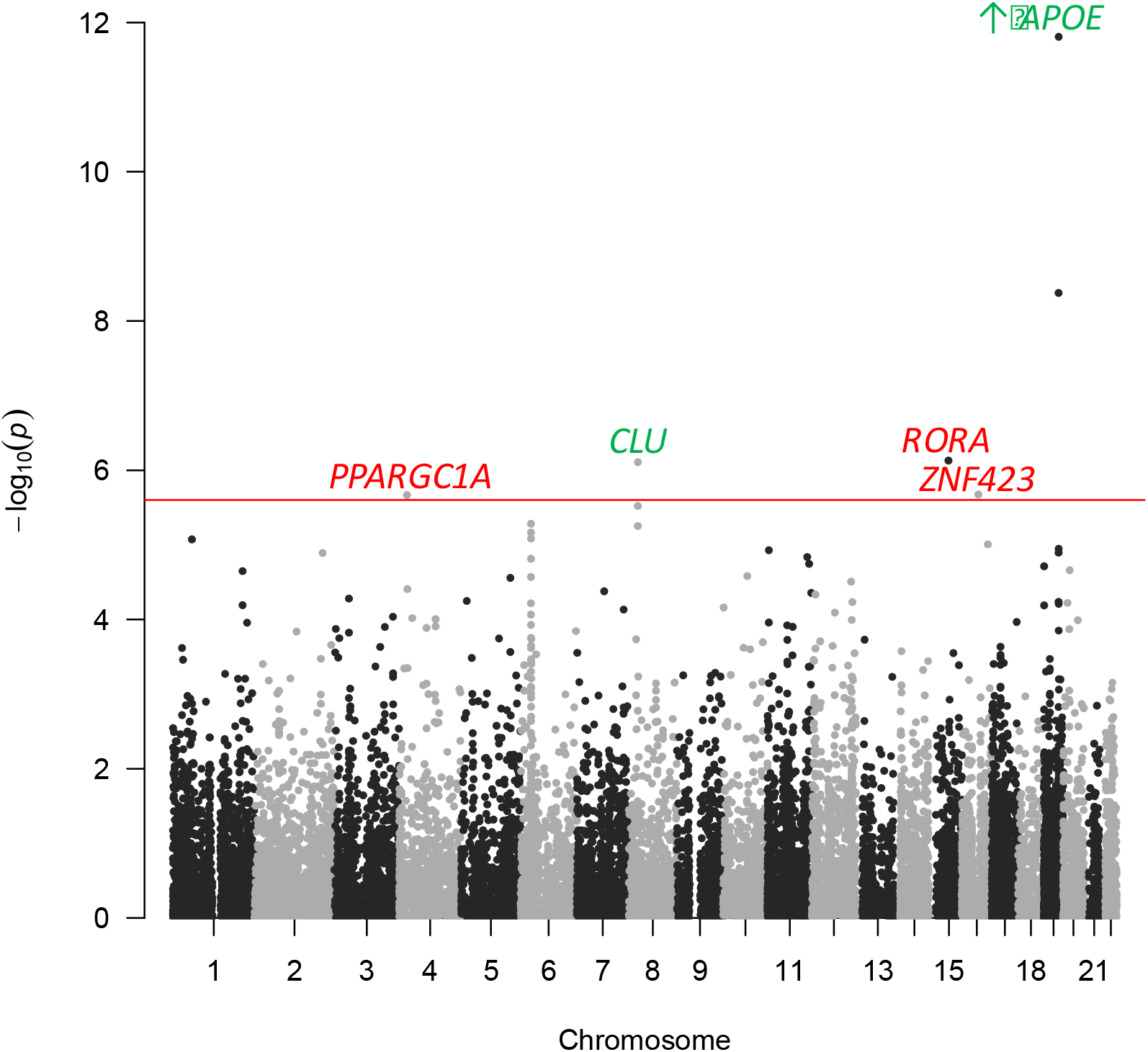
Manhattan Plot for the POLARIS Gene-Based Analysis in Imputed GERAD Data Using a Gene Window 35kb Upstream and 10kb Downstream

The gene expression patterns in these novel genes were investigated using the BRAINEAC (Trabzuni et al. [2011]) database from the UK Brain Expression Consortium. For the *PPARGC1A* gene, the SNP rs67436520 has the best cis-expression quantitative trait loci (eQTL) p-value of 3.3×10^-4^, this is expressed in the hippocampus. The best cis-eQTL p-value in the *RORA* gene is 1.5×10^-4^, this is for SNP rs113223478 which is 78.5kb upstream of the gene, and is expressed in the substantia nigra. Finally, SNP rs2270396 has the best cis-eQTL p-value in the *ZNF423* gene with a p-value of 3.0×10^-5^ and is expressed in the frontal cortex.

The gene-based analysis results from MAGMA-PCA and MAGMA-SUMMARY using a gene window identified genes which are in the *APOE* region (results are not shown).

## Conserved Regions

GWAS data was interrogated for genes that are evolutionarily constrained, since such constraint indicates functional importance (Babarinde et al. [2016]). Genes from the gene-based analysis which are in the Exome Aggregation Consortium database (Lek et al. [2016]) and LoF intolerant were determined. Additionally, genes which overlap with conserved noncoding sequences were found (Babarinde et al. [2016]). It was then assessed whether there was enrichment for LoF intolerant genes or genes in conserved noncoding sequences using the POLARIS gene-based results in imputed data. There are 557 independent genes which do not overlap, for these genes, there is no enrichment for genes in conserved regions at either the nominal or gene-wide p-value threshold (OR=0.95, p=0.9052 and OR=4.19, p=0.3505 respectively). Similarly, no evidence of an enrichment of genes in conserved noncoding regions at either the nominal or gene-wide p-value threshold (OR=2.97, p=0.5586 and OR=31.57, p=1 respectively). Similar results were obtained for MAGMA derived p-values (results are not shown).

In summary, there is consistently no enrichment at the gene-wide p-value threshold for genes in either loss of function intolerant or conserved noncoding sequence regions, potentially due to the fact that AD is a post-reproductive disorder.

## Discussion

A gene-based analysis was performed using the individual genotypes in the GERAD imputed data and the summary statistics from IGAP data excluding the GERAD subjects was used to inform the analysis. When no gene window is considered, POLARIS does not identify any novel genes, only genes on chromosome 19 which are likely influenced by the large effect of *APOE*. When a gene window was expanded around the gene, 35kb upstream and 10kb downstream, three novel genes were found from the POLARIS gene-based analysis. The gene window is likely to include transcriptional regulatory elements in the gene (Network and Consortium Pathway Analysis Subgroup of Psychiatric Genomics [2015]) and thus contain SNPs influencing gene expression.

Three novel genes were found to be associated with AD using the POLARIS method only. Gene-based analyses with MAGMA software identified only genome-wide significant genes in the *APOE* region. POLARIS determines a larger number of gene-wide significant genes compared to both the MAGMA approaches. This is consistent when the significance threshold is altered based on the number of genes from the analysis (0.05/no. genes). Supplementary Figures S1 and S2 show the gene p-value comparison between POLARIS and MAGMA-PCA and MAGMA-SUMM respectively; POLARIS generally has more significant p-values compared to both MAGMA approaches. This is likely due to the fact that MAGMA uses the GERAD data only whereas POLARIS additionally informs the analysis with the IGAPnoGERAD data. Thus, increasing power while maintaining the independence of the GERAD dataset. The novel genes are *PPARGC1A, RORA* and *ZNF423*, all of which have biological relevance to AD.

*PPARGC1A* (peroxisome proliferator-activated receptor gamma co-activator 1alpha) is a transcriptional coactivator involved in a wide range of cellular and physiological functions. It is part of the PGC-1 family of transcriptional coactivators that mainly regulate mitochondrial biogenesis to in turn regulate the cellular energy metabolism (Luo et al. [2016]). The gene product, PGC-1α, is an interacting partner of a wide range of nuclear receptors and transcription factors. It is associated with a wide-range of biological processes (Gene Ontology, http://www.ensembl.org), including response to a variety of cellular and external stimuli, cellular glucose homeostasis, circadian rhythm, regulation of neuron apoptosis, etc. Previous animal model work has shown that overexpression of hPGC-1α in APP23 mice improved spatial and recognition memory, along with significant reduction of Aβ deposition (Katsouri et al. [2016]). Furthermore, hPGC-1α overexpression also reduced the levels of proinflammatory cytokines and microglial activation (Nijland et al. [2014], Katsouri et al. [2016]). This suggests a direct link with recent genetic evidence of microglia-mediated innate immune response involvement in Alzheimer’s Disease (Sims et al. [2017]). In addition, an activation of PGC-1α by EKR and p38 inhibitors have been shown to improve spatial and learning memory in Aβ-injected rats (Ashabi et al. [2012]). *PPARGC1A* has also been implicated in the pathogenesis of other neurodegenerative disorders, namely Huntington’s and Parkinson’s diseases (Tsunemi et al. [2012]).

Retinoic acid receptor-related orphan receptor alpha (*RORA*) is a nuclear hormone receptor and is involved in a variety of functions, circadian rhythm, cholesterol metabolism, inflammation, etc (Jetten et al. [2009]). *RORA* binds to genomic regions of transcription start sites of more than 3,000 genes in human monocytic and endothelial cell lines (Gulec et al. [2017]). *RORA* and *PPARGC1A* are close biological partners, with PGC-1α stimulates the expression of a number of clock genes through the coactivation of the ROR family of orphan nuclear receptors (Liu et al. [2007]). *RORA* has been shown to be linked to other genes previously implicated in AD (Hu et al. [2017]) and has also been implicated in a large number of neuropsychiatric disorders, such as post-traumatic stress disorder (Miller et al. [2013]). Furthermore, *RORA* trans-activates Il-6 and is thought to be neuro-protective in astrocytes and anti-inflammatory in peripheral tissues (Journiac et al. [2009]). The two genes, *RORA* and *PPARGC1A* that we report here provide further evidence of the involvement of inflammation in the pathogenesis of Alzheimer’s disease.

Finally, *ZNF423* is a nuclear protein that belongs to the Kruppel-like C2H2 zinc finger proteins. *ZNF423* directs bone morphogenetic protein (BMP)-dependent signalling activity and aberrant forms impede B cell differentiation (Harder et al. [2013]). Furthermore, an increased gene-expression of *ZNF423* has been associated in patients with systemic lupus erythematosus pointing to an impaired function of B cells in human mesenchymal stem cells (Feng et al. [2014]). *ZNF423* resides in an AD-specific protein network (Hu et al. [2017]). *ZNF423* is likely involved in DNA damage repair (Chaki et al, [2012]). Previously, it also has been shown that missense and LoF variants are likely pathogenic for abnormality of brain morphology, Joubert syndrome and Nephronophthisis with autosomal dominant or autosomal recessive inheritance (www.omim.org, https://www.ncbi.nlm.nih.gov/clinvar/). These disorders present with a range of phenotypic characteristics, with the central nervous system being affected too (more specifically the cerebellar vermis). In nur12 mouse model (with introduced nonsense mutation in exon 4 of the mouse Zfp423 gene), Alcaraz et al. [2006] observed loss of the corpus callosum, reduction of hippocampus, and a malformation of the cerebellum reminiscent of patients with Dandy-Walker syndrome. Within the cerebellum, Zfp423 was observed to be expressed in both ventricular and external germinal zones. Loss of Zfp423 was also observed to lead to diminished proliferation by granule cell precursors in the external germinal layer and abnormal differentiation and migration of ventricular zone-derived neurons and Bergmann glia (Alcaraz et al. [2006]).

Genes determined from the gene-based analysis show no enrichment in conserved regions, in either evolutionary constrained LoF intolerant regions or in conserved noncoding sequences. This is expected in AD because it is a post-reproductive disorder. LoF highly penetrant rare variants in variation intolerant genes have been strongly linked to neurodevelopmental disorders with paediatric onset previously (Grozeva et al. [2015], Deciphering Developmental disorders study [2017]). Highly penetrant rare variants have been linked to familial AD with an early onset, but for the majority of individuals developing AD, this is not the case. In addition, rare copy number variants affecting haploinsufficient genes have been linked to SZ, but not AD (Grozeva et al. [2012], Chapman et al. [2013]). Therefore, it could be suggested that the mechanism of disease leading to AD, is different to mechanism leading to diseases with paediatric onset and are of neurodevelopmental origin. This is pointing to divergent biological mechanisms across the common and rare variant signals and between disorders polygenic in nature.

## Appendix 1

Authors who contributed to the generation of original study data for GERAD, ADGC, CHARGE and EADI, but not to the current publication are included herein. Author affiliations can be found in the Supplementary material.

## Acknowledgements

We thank the MRC Centre for Neuropsychiatric Genetics and Genomics for supporting this project and the MRC for supporting author Emily Baker. This project was also supported by Dementia Project UK (DPUK). Data used in the preparation of this article were obtained from the Genetic and Environmental Risk for Alzheimer’s disease (GERAD1) Consortium (Harold et al. [2009]). We also thank the Dementia Research Institute (DRI) for supporting this project.

We thank the International Genomics of Alzheimer’s Project (IGAP) for providing summary results data for these analyses. The investigators within IGAP contributed to the design and implementation of IGAP and/or provided data but did not participate in analysis or writing of this report.

## Supplementary Material

**Figure S1:**
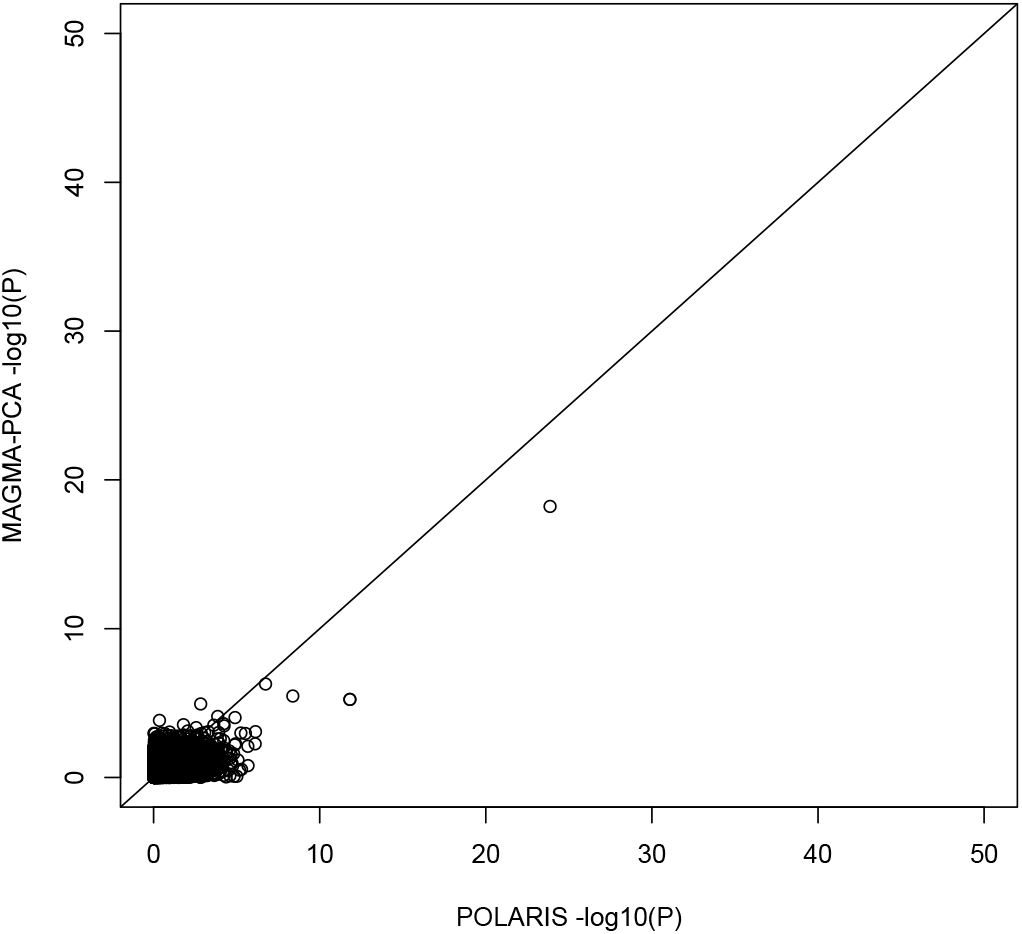
Gene -log_10_(p-value) Comparison Between POLARIS and MAGMA-PCA, with gene window

**Figure S2:**
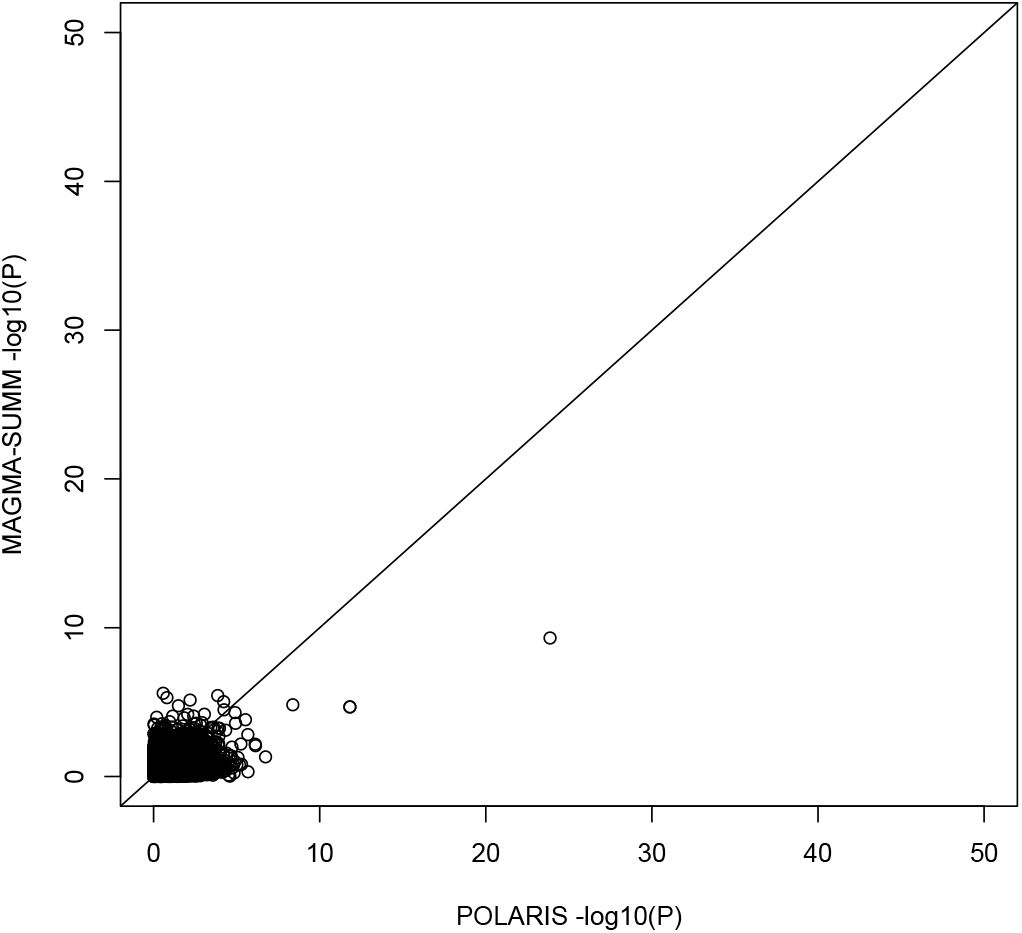
Gene -log_10_(p-value) Comparison Between POLARIS and MAGMA-SUMMARY, with gene window

**Figure S3:**
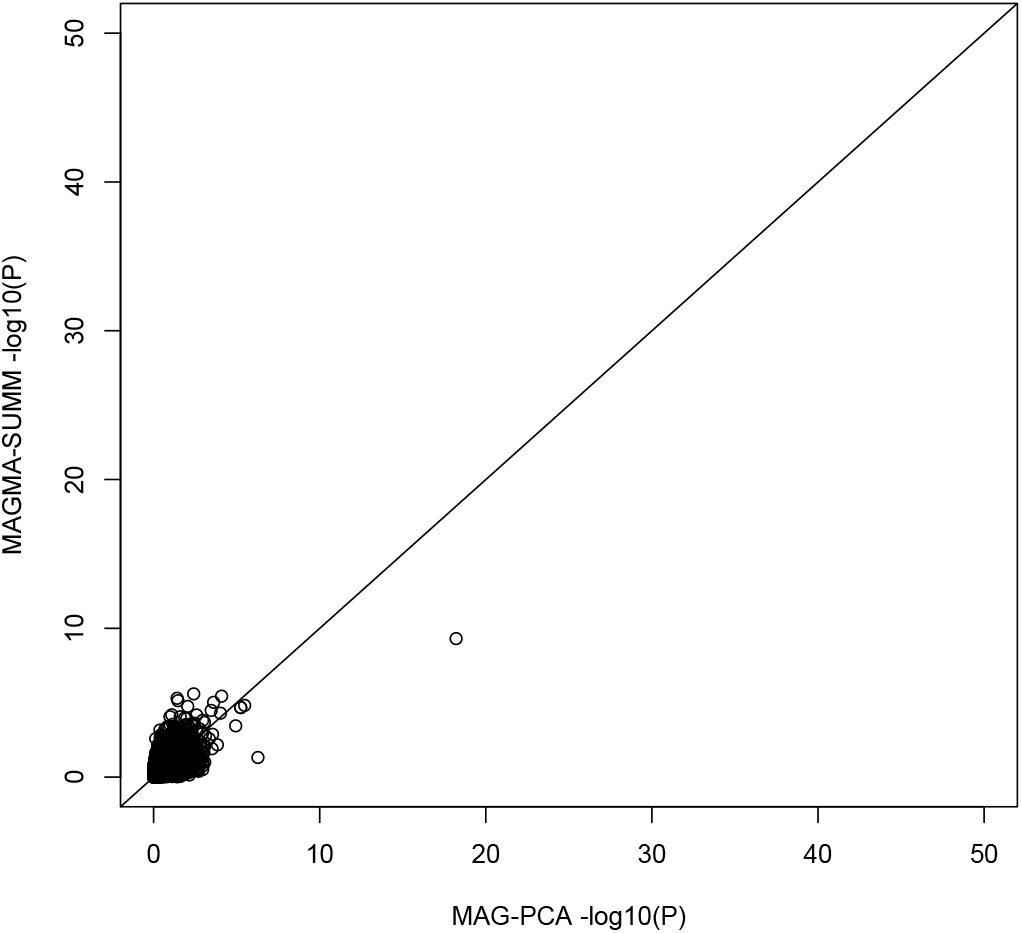
Gene -log_10_(p-value) Comparison Between MAGMA-PCA and MAGMA-SUMMARY, with gene window

